# Homologous laminar organization of the mouse and human subiculum

**DOI:** 10.1101/2019.12.20.883074

**Authors:** Michael S. Bienkowski, Farshid Sepehrband, Nyoman D. Kurniawan, Jim Stanis, Laura Korobkova, Neda Khanjani, Houri Hintiryan, Carol A. Miller, Hong-Wei Dong

## Abstract

The subiculum is the major output structure of the hippocampal formation and one of the brain regions most affected by Alzheimer’s disease. Our previous work revealed a hidden laminar architecture within the mouse subiculum. However, the rotation of the hippocampal longitudinal axis across species makes it unclear how the laminar organization is represented in human subiculum. Using *in situ* hybridization data from the Allen Human Brain Atlas, we demonstrate that the human subiculum also contains complementary laminar gene expression patterns similar to the mouse. In addition, we provide evidence that the molecular domain boundaries in human subiculum correspond to microstructural differences observed in high resolution MRI and fiber density imaging. Finally, we show both similarities and differences in the gene expression profile of subiculum pyramidal cells within homologous lamina. Overall, we present a new 3D model of the anatomical organization of human subiculum and its evolution from the mouse.

## Introduction

The subiculum (SUB) is a stratified cortical region that is anatomically-positioned as the major output of the hippocampal formation. The SUB strata consist of a plexiform molecular layer, a pyramidal cell layer, and a deep polymorphic cell layer ^1^. Based on Golgi stained neuronal morphology, Lorente de Nó described an additional sublaminar organization within the pyramidal layer that he used to define SUB subfields including the prosubiculum (ProSUB) ^2^. Although Lorente de Nó could discern these more subtle SUB cellular lamina across multiple mammalian species, mapping the complete laminar distribution of SUB pyramidal neurons and clearly identifying SUB subfield organization has remained challenging. Later, separate anatomical tract tracing studies in rats suggested both columnar and laminar organization characterized SUB neuron connectivity ^3–5^, but a comprehensive understanding as to how many lamina and columns existed along the whole longitudinal axis remained obscure until recently.

Our previous work creating the Hippocampus Gene Expression Atlas (HGEA) demonstrated that combinatorial gene expression patterns identify the hidden sublaminar organization of SUB pyramidal neurons and these gene expression patterns were highly related to specific connectivity labeling patterns ^6^. Unlike anatomical tracer patterns which typically label a topographic subpopulation defined by the size and placement of the injection site, *in situ* hybridization gene expression patterns reveal a complete laminar distribution across the entire longitudinal axis. Similar to the approach used by Lorente de Nó previously, the HGEA outlines five SUB subregions based on the representation of four identified gene expression lamina: dorsal and ventral parts of the dorsal subiculum (SUBdd and SUBdv, respectively), the prosubiculum (ProSUB), and the ventral subiculum (SUBv) along with its ventral tip (SUBvv). While the HGEA delineates the complete distribution of SUB subregion and lamina across the longitudinal axis in mice, it remains unclear how the new HGEA subregional and laminar organization is represented across other mammals, particularly humans. Previous translational studies have examined gene expression patterns to define hippocampal and SUB boundaries, but did not report a laminar organization within the SUB ^7–9^.

Many functional and anatomical studies have suggested that the hippocampus is generally homologous across mammals although the spatial position of the hippocampus within the brain has shifted across evolution ^3,10–17^. The hippocampal longitudinal axis (red axis in **Fig. 1**) is oriented dorsoventrally in mice, whereas in primates and humans this axis is rotated into the posterior-anterior direction. Based on this rotation, the mouse dorsal SUB is generally believed to be homologous to the human posterior SUB and mouse ventral SUB is homologous to anterior human SUB. Functional evidence supports this view as the mouse dorsal SUB and human posterior SUB are involved in visuospatial navigation ^10,17–19^. In contrast, the mouse ventral SUB and human anterior SUB are related to limbic emotional processing and social behaviors ^12,20,21^. Based on the anatomical and functional homology, we hypothesized that the laminar gene expression patterns that delineate mouse HGEA SUB subregions would demonstrate a similar relationship pattern in the corresponding parts of human SUB. Analyzing gene expression patterns from the Allen Human Brain Atlas *in situ* hybridization database, we identified genes with unique spatial distribution patterns that characterize relatively distinct and complementary lamina across the human posterior and anterior SUB. Particularly, the posterior human SUB contains three complementary gene expression layers whose distribution patterns strongly reflect the mouse SUBdd and ProSUB when accounting for the rotation of the longitudinal axis. Additionally, posterior SUB gene expression boundaries aligned with differences in fiber density microstructure demonstrated by *ex vivo* diffusion-weighted magnetic resonance imaging (DW-MRI). In the anterior SUB, we found these lamina continued within ventral parts of the SUB, but not in the dorsal parts of the anterior SUB, suggesting a difference between dorsal and ventral parts of the anterior SUB that was not previously known. Taking this new data together with our previous understanding of the mouse subiculum, we propose a new 3D model of the human subiculum and its homology to the rodent. Overall, our gene expression analysis provides a new understanding of the homology between mouse and human hippocampus that is critical to translational studies of hippocampal diseases such as Alzheimer’s disease, hippocampal sclerosis, and epilepsy.

**Figure. 1.**
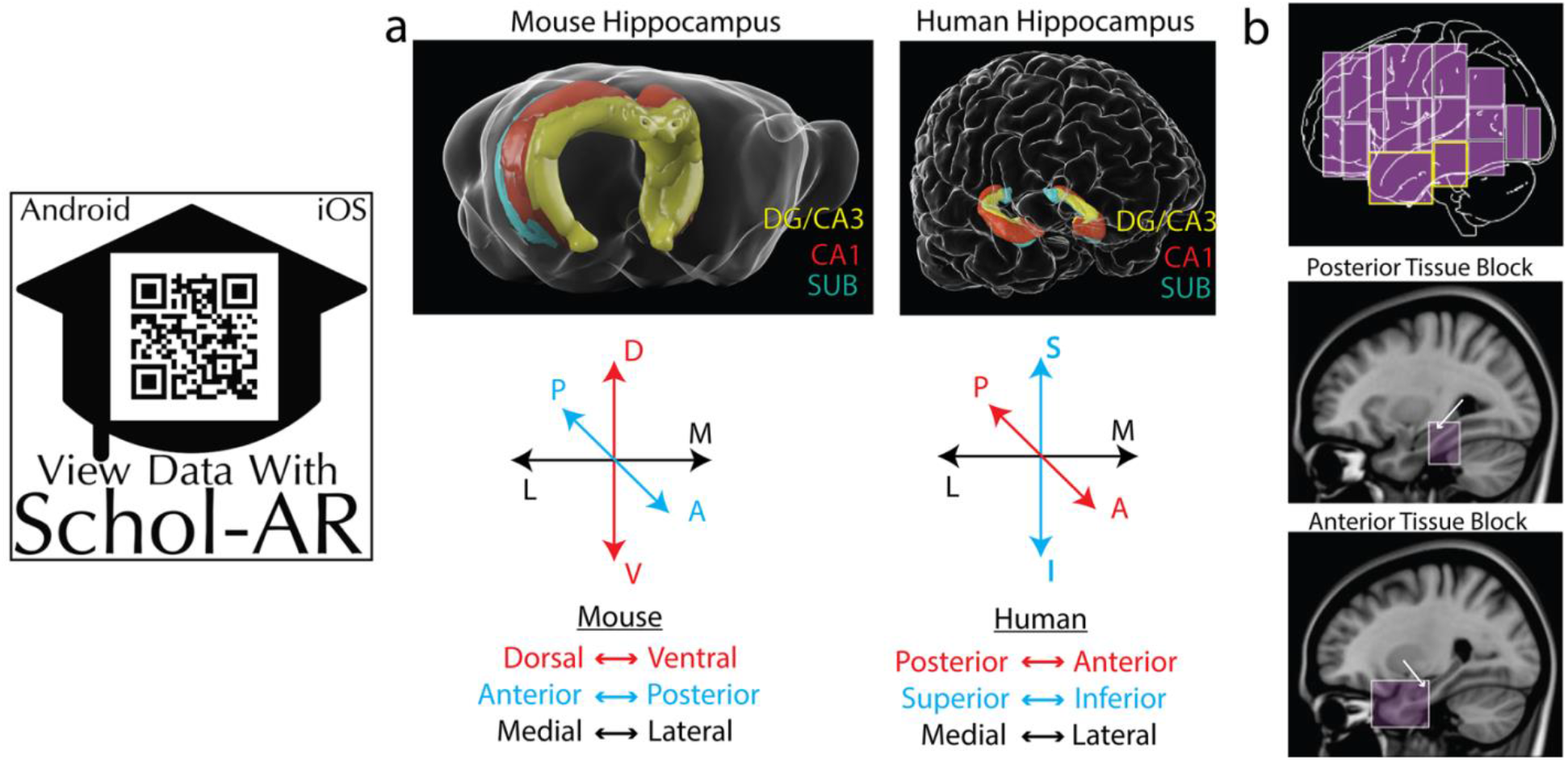
Structural comparison of the mouse and human hippocampus. **a** Three-dimensional representations of the mouse and human hippocampus showing the relative locations of the DG/CA3 (yellow), CA1 (red), and SUB (green); see also Supplementary Movie 1 or use Schol-AR app). The longitudinal hippocampal axis (red in axes chart) in mouse is oriented dorsoventrally, whereas the human longitudinal axis is rotated into the anterior-posterior axis. In addition the anterior/posterior (septo-temporal; blue color) axis in mice is oriented in the superior-inferior direction in humans. **b** Sagittal view of human brain volume representing the spatial location of all *in situ* hybridization datasets from the Allen Human Brian Atlas (top). Two tissue blocks containing posterior (middle) and anterior (bottom) parts of the hippocampus (white arrows) are shown overlaid on sagittal 3T structural MRI images. All images in **b** are from www.brain-map.org. Data viewable with Schol-AR augmented reality app, for details visit https://www.ini.usc.edu/scholar/download.html.

## Results

All *in situ* hybridization data was downloaded from the online Allen Brain Atlas image database (www.brain-map.org). Compared to the mouse database which includes both coronal and sagittal tissue sections, the current Allen Human Brain Atlas database contains only a limited set of *in situ* hybridization data in coronally-sectioned tissue (www.human.brain-map.org). Two separate tissue blocks containing hippocampus/amygdala were found in four subjects as part of the Neurotransmitter Study: one block of tissue includes the anterior hippocampal pole and another block of tissue at a more intermediate/posterior hippocampal level. We observed relatively consistent gene expression patterns in all 4 cases although because of variability in tissue dissection and sectioning quality, it was difficult to relate corresponding sections across subjects. Therefore, we present data from the tissue series with the best histological quality (H0351.1010, 28 year old Hispanic male with no known cognitive impairment) to demonstrate their patterns across adjacent rostrocaudal sections (tissue index (TI) identifies adjacent series section number). We will first describe observed gene expression patterns in the human posterior SUB followed by analysis of anterior SUB gene expression patterns. In addition, we provide evidence from *ex vivo* MRI imaging data that gene expression boundary delineations correspond to differences in human imaging microstructure. Finally, we provide a comparison of mouse and human gene expression within homologous SUB lamina.

### Gene expression patterns in the posterior human SUB

Within posterior hippocampal tissue sections, *in situ* hybridization gene expression patterns reveal the laminar organization of SUB pyramidal neurons. Notably, neurons expressing *Nts*, *Chrm2*, and *Htr2a* are broadly distributed as three complementary gene expression patterns across the CA1, ProSUB, and SUB (**Fig. 2**). *Nts* is robustly expressed within a layer of superficial SUB pyramidal neurons (adjacent to the molecular layer), whereas *Chrm2*-expressing pyramidal neurons are distributed as a deep thin layer directly adjacent to the alveus white matter tract. The deep *Chrm2* gene expression layer is thicker laterally where Chrm2 expression also continues into the presubiculum (PRE), but gradually becomes thinner as it extends medially into the ProSUB region. In contrast to the SUB, the superficial ProSUB neurons near the molecular layer robustly express *Htr2a* rather than *Nts*. *Htr2a* expression is continuous within CA1 and ProSUB, but ends near the SUB border. Together, *Nts*, *Chrm2*, and *Htr2a* expression patterns identify three separate gene expression domains across the human SUB and ProSUB with relatively distinct boundaries. The SUB and ProSUB can be distinguished by *Nts* vs. *Htr2a* expression although each region contains a common deep layer of *Chrm2*-expressing neurons.

**Figure. 2.**
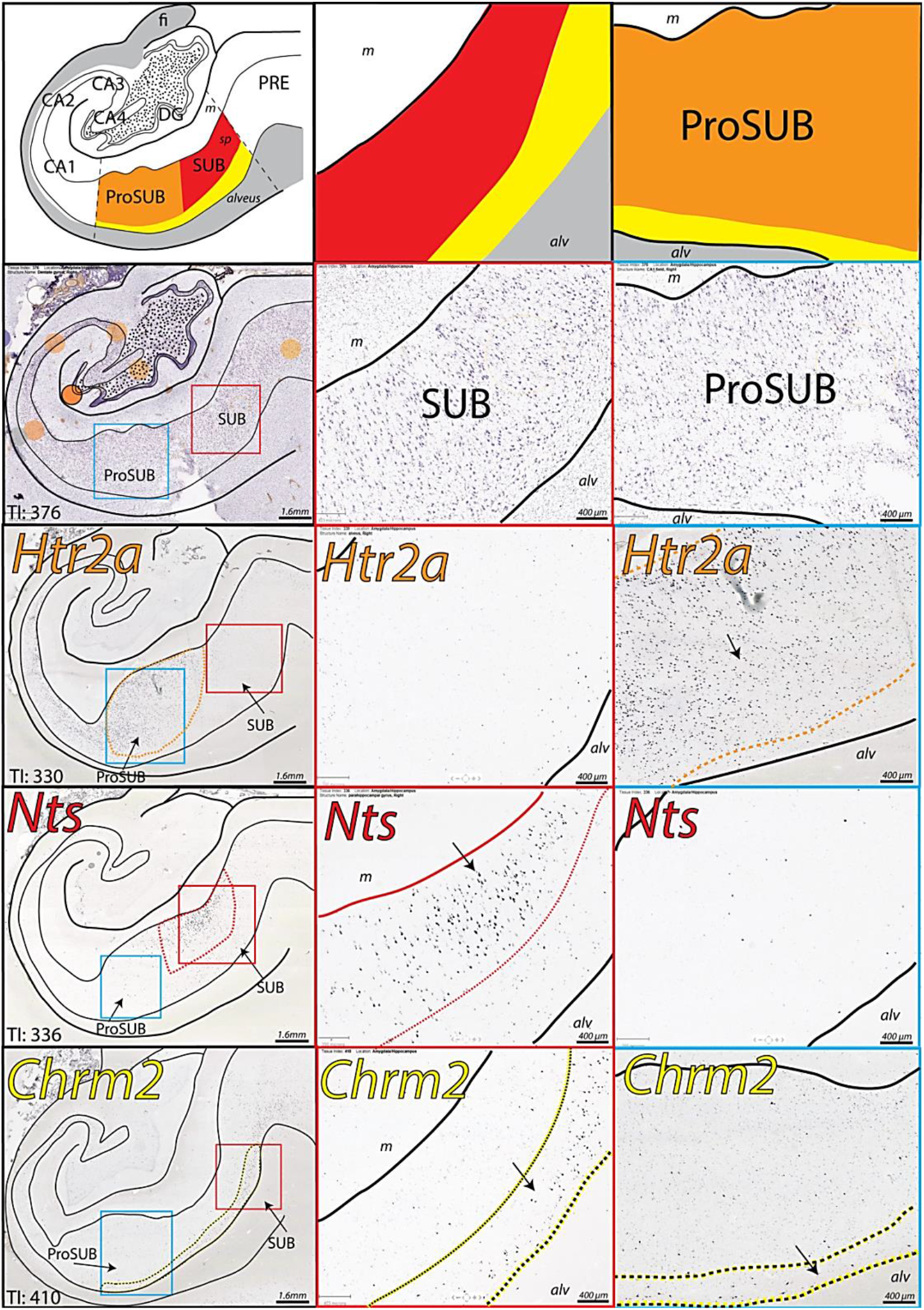
Complementary gene expression patterns in the posterior subiculum. (left) *In situ* hybridization staining for *Htr2a*, *Nts*, and *Chrm2* in adjacent tissue sections alongside Nissl stained cytoarchitecture and corresponding atlas drawing (based on Nissl section). *Htr2a* is strongly expressed in the CA1 and superficial ProSUB area (outlined in orange), *Nts* is strongly expressed in superficial SUB cells located near the molecular layer (m; expression area outlined in red), and *Chrm2* is expressed in a deep layer of cells dorsal to the alveus (alv) within the ProSUB and SUB (outlined in yellow). Tissue Index (TI) number references section number within the overall tissue series. Red and blue boxes represent zoomed in image areas of the SUB and ProSUB, respectively, shown in middle and right columns. Together, the three gene expression patterns represent disparate molecular domains as shown by the red, orange, and yellow colored regions in the atlas drawings in top row. All Nissl and *in situ* hybridization images downloaded from www.brain-map.org.

Along the longitudinal axis, the size and shape of the three SUB/ProSUB gene expression domains and their boundaries shift as the cytoarchitecture of the SUB and ProSUB changes (**Fig. 3**). However, the complementary arrangement of the gene expression domains to each other, as well as their relative position adjacent to the alveus and molecular layer, remains consistent. From the available consecutive tissue series covering 10mm of the hippocampal longitudinal axis, the data suggests that the relatively segregated gene expression layers can be considered as continuous laminar sheets extending rostrocaudally across the whole SUB. Additionally, gene expression for *TH* and *PCP4* appear to mirror the distribution patterns of *Nts* and *Chrm2*, respectively (**Fig. 3**). Although it is unclear if these genes are expressed within the same individual cell types or shar a similar spatial distribution, *TH* and *PCP4* gene expression demonstrates that the SUB molecular domains represent differences in multiple combinatorial gene expression patterns.

**Figure. 3.**
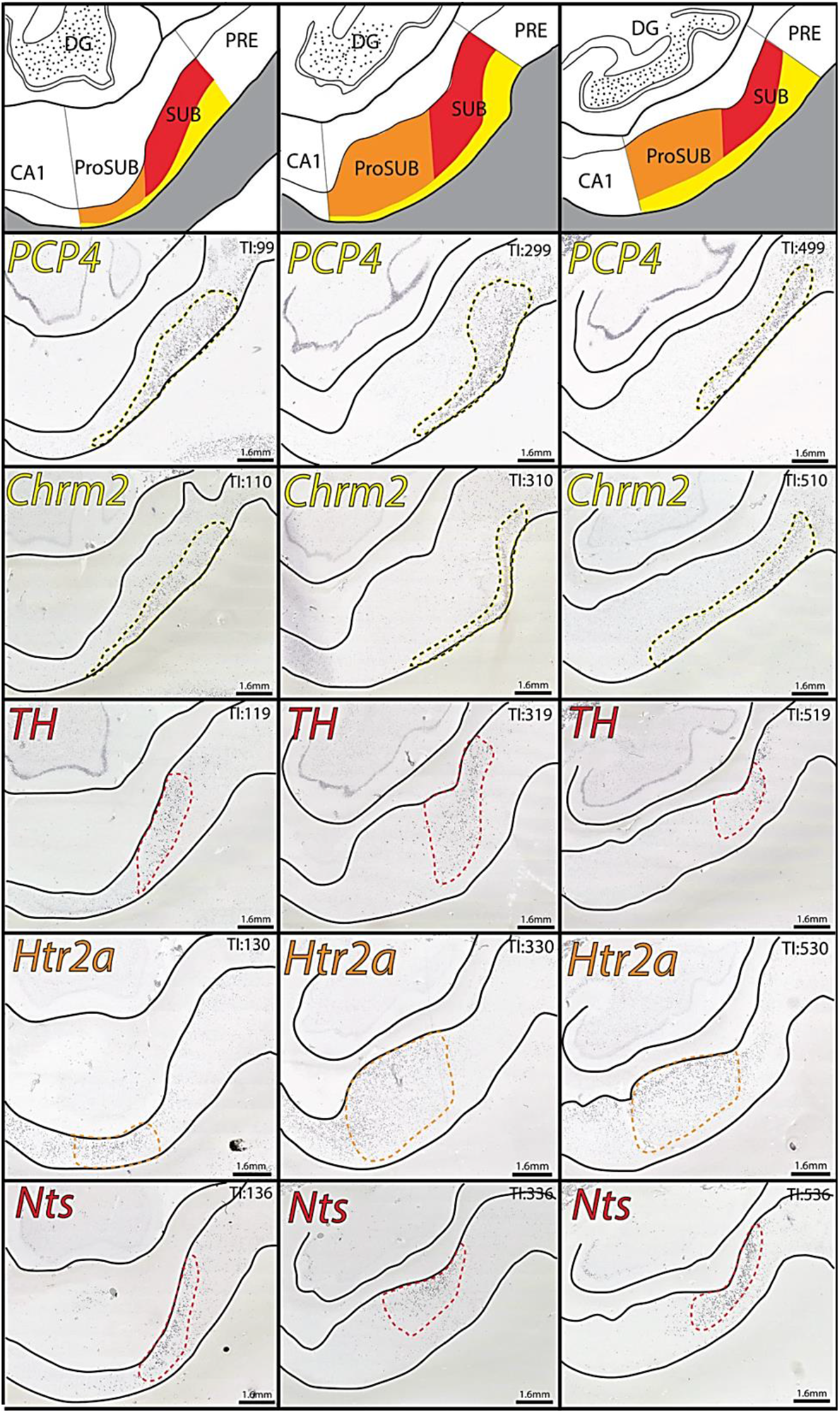
Gene expression domain boundaries shift across the longitudinal axis, but maintain their complementary relationship. *In situ* hybridization images of distribution patterns for *PCP4*, *Chrm2*, *TH*, *Htr2a*, and *Nts* gene expression (outlined in yellow, red, or orange colors) at three different rostrocaudal levels spanning 8mm of the longitudinal axis (posterior to anterior from left to right). Tissue Index (TI) numbers on each image reference section number in tissue series. *PCP4* and *TH* expression patterns closely mirror the distribution patterns of *Chrm2* and *Nts*, respectively. On the top are corresponding atlas drawings demarcating each of the three laminar molecular domains (drawings based on the boundaries drawn in the *TH*-labeled sections). Note, at most posterior levels (TI: 99-136), the area of the ProSUB is small compared to the more anterior levels. All *in situ* hybridization images downloaded from www.brain-map.org.

### Gene expression patterns in the anterior human SUB

Compared to the posterior hippocampus, the anterior hippocampus is structurally more complex including cytoarchitectural ridges and folding. At the anterior pole of the hippocampus, the transverse axis folds around the dentate gyrus and ultimately ends near the amygdala, an area commonly referred to as the hippocampal amygdala transition area (HATA). In coronal sections of the anterior hippocampus, the CA1/SUB is present on both dorsal and ventral sides of the hippocampal sulcus. In the ventral part of anterior SUB (**Fig. 4**), *in situ* hybridization reveals that *Nts*-, *Htr2a*-, and *Chrm2*-expressing neurons are arranged in similar gene expression patterns to the posterior SUB levels, suggesting the ventral anterior SUB neurons are rostrocaudally continuous with neurons observed in the posterior SUB sections (**Fig. 3**). In contrast, the dorsal part of anterior SUB demonstrates a distinct combination of gene expression and cytoarchitecture. Examining the Nissl cytoarchitecture of this region reveals a distinct intermediate layer of densely packed and darkly stained pyramidal neurons that suggests a trilaminar organization (**Fig. 4**). *Chrm2*-expressing neurons are distributed in the deep layer adjacent to the white matter whereas *Htr2a*-expressing neurons form a complementary gene expression pattern and are located within both the intermediate and superficial layer (*Nts* expression is mostly absent).

**Figure. 4.**
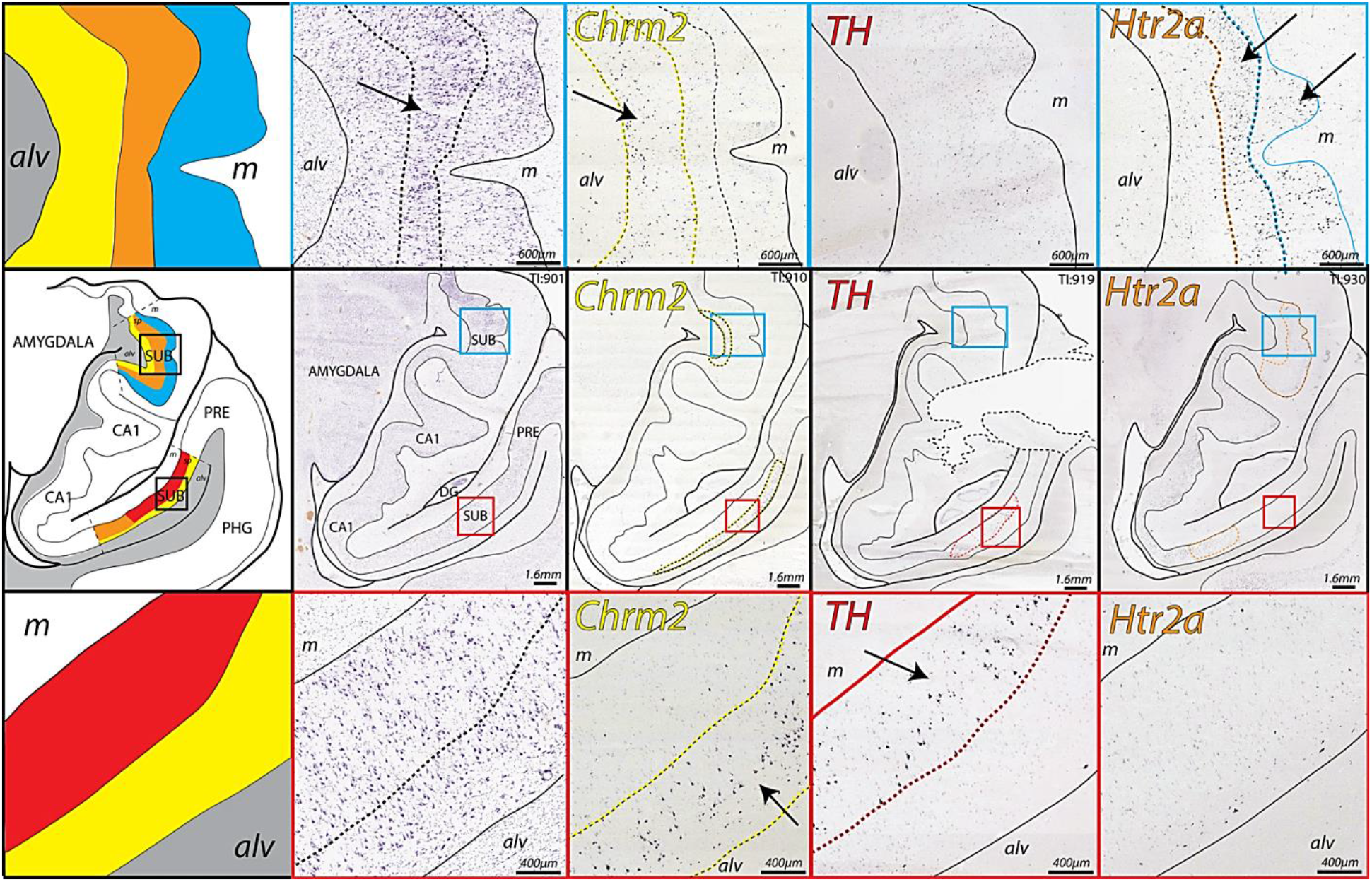
Complementary gene expression patterns in the anterior SUB (TI: 901-930). (middle row) *In situ* hybridization staining for *Chrm2*, *TH*, and *Htr2a* (colored outlined) in adjacent tissue sections alongside Nissl stained cytoarchitecture and corresponding atlas drawing (based on Nissl section). In anterior coronal sections, the SUB appears separated by the CA1 into a dorsal and ventral region. The ventral region of the SUB contains complementary *Chrm2*, *TH*, and *Htr2a* expression patterns in the SUB and ProSUB as observed in posterior hippocampal levels (zoomed in images of red boxed regions are shown in bottom row). In contrast, the dorsal region of the SUB at this anterior level contains *Chrm2* and *Htr2a* expression, but very little *TH* expression (zoomed in images in blue boxed regions are shown in top row). Closer examination of the Nissl staining suggests a tri-laminar cytoarchitecture containing a distinct intermediate layer (arrow in top row Nissl image, orange domain in atlas drawing) with cells that are more darkly-stained and densely packed than the cells located deeper (yellow in atlas drawing) and more superficially (blue in atlas drawing). Based on this trilaminar organization, Chrm2-expressing cells are primarily distributed in the deep layer adjacent to the alveus (alv) whereas *Htr2a*-expressing cells are located in the intermediate and superficial layers near the molecular layer (m). All Nissl and *in situ* hybridization images downloaded from www.brain-map.org.

Examining tissue sections closer to the anterior pole reveals that the dorsal and ventral parts of the SUB are continuously adjoined. The layer of *Chrm2*-expressing neurons in the dorsal part of anterior SUB joins with the ventral part of SUB by continuing around the medial/anterior pole of the hippocampus (**Fig. 5**). In addition, the *Nts*-expressing neuronal layer also partially extends around the medial/anterior pole where these cells seem to be interposed between the layers of *Chrm*- and *Htr2a*-expressing neurons. At the anterior end of the SUB, *Chrm2*-expressing neurons are distributed as an outer ring layer with a more internal core of *Nts*-expressing neurons and a relative absence of *Htr2a*-expressing neurons. Overall, the distribution of gene expression patterns suggests that the SUB and ProSUB contain unique combinations of molecular domains composed of continuous sheets of distinct neuronal cell types spanning the entire hippocampal axis.

**Figure. 5.**
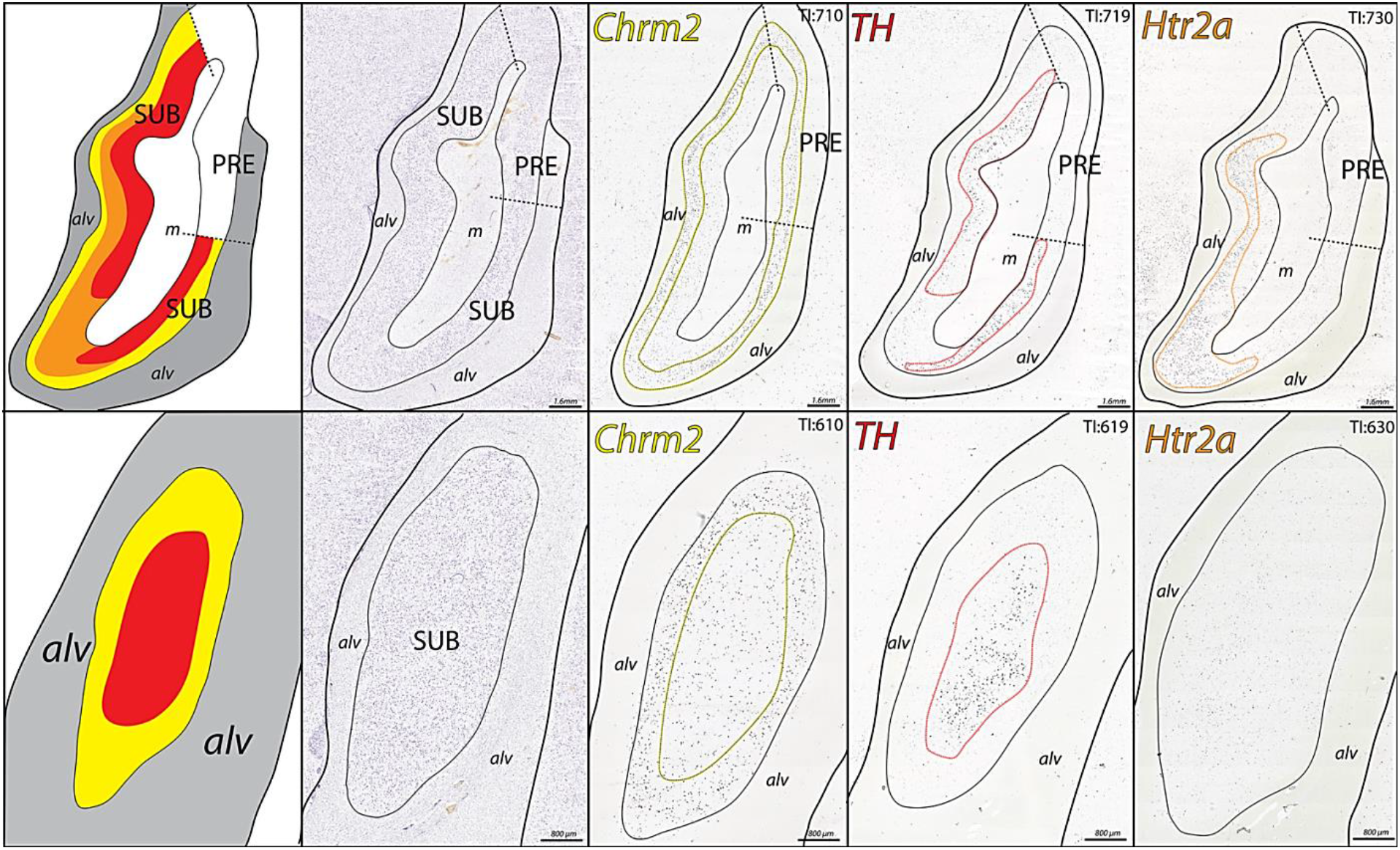
Gene expression patterns at the SUB anterior tip. (top row) 4mm anterior to the section shown in Fig. 4, the CA1 is no longer present and the ventral and dorsal regions of the SUB become continuous along with their laminar gene expression patterns. *In situ* hybridization images of *Chrm2*, *TH*, and *Htr2a* expression (colored outlines) are presented with adjacent Nissl and corresponding atlas drawings. *Chrm2*-expressing cells form a continuous external layer adjacent to the alveus (alv), whereas *TH* and *Htr2a* expression is distributed within cells located more internally near the molecular layer (m). Unlike the tissue level shown in Fig. 4, there is abundant TH expression in the more dorsal part of the SUB. In the most anterior coronal section (bottom row, 2mm anterior to sections shown in top row), the molecular layer is no longer present. *Chrm2*-expressing cells form a continuous ring layer that surrounds a core of TH expressing cells (few *Htr2a* expressing cells remain at this level). Together this data shows that *Chrm2*, *TH*, and *Htr2a* expressing cells form three distinct layers that wrap continuously around the anterior/medial tip of the hippocampus. In relation to the section in Fig. 4, the layer of *TH* expression appears to end dorsally approximately ~6mm from the anterior pole of the SUB, whereas *Chrm2* and *Htr2a* expression continues posteriorly within the dorsal part of the anterior SUB.). All Nissl and *in situ* hybridization images downloaded from www.brain-map.org.

### Comparison of gene expression-derived histological boundaries to human MRI imaging

To determine if the regional boundaries identified by different gene expression patterns corresponded to other structural differences in connectivity microstructure, we compared our gene expression delineations to *in vivo* and *ex vivo* structural MRI and track density imaging (TDI) of hippocampal pathways (**Fig. 6**). At a rostrocaudal level similar to the posterior SUB described above, *in vivo* 7T MRI images of human brain provide macroscopic resolution, sufficient to clearly distinguish the SUB molecular layer from the pyramidal layer, but difficult to distinguish finer details that are useful for delineating subregion boundaries (**Fig. 6a,b**). In comparison, *ex vivo* 16.4T MRI of a dissected human hippocampal sample provides mesoscopic details. Hippocampal strata are clearly apparent, including the granule cell layer of the dentate gyrus, and the alveus can be distinguished from the angular bundle within the white matter (**Fig 6c**). TDI analysis of the tissue microstructure in this sample reveals differences in the fiber pathway orientation and distribution (**Fig. 6d,e**) that notably align to our observed gene expression boundaries (**Fig. 6f**). The PRE contains dense bundles of dorsoventrally-oriented perforant path fibers that strongly contrast with more minimal fibers within the SUB. Within the boundaries of the ProSUB, thicker dorsoventally oriented fiber bundles are apparent that contrast it from the adjacent CA1 and SUB. Finally, a mediolateral fiber bundle can be observed running along the deep ProSUB and SUB just dorsal to the alveus that appears similar to the laminar distribution of *Chrm2*-expressing neurons. Overall, the TDI fiber analysis provides evidence that microstructural differences in human hippocampal tissue correspond to differences in genetic cell type distribution when observed with high enough resolution.

**Figure. 6.**
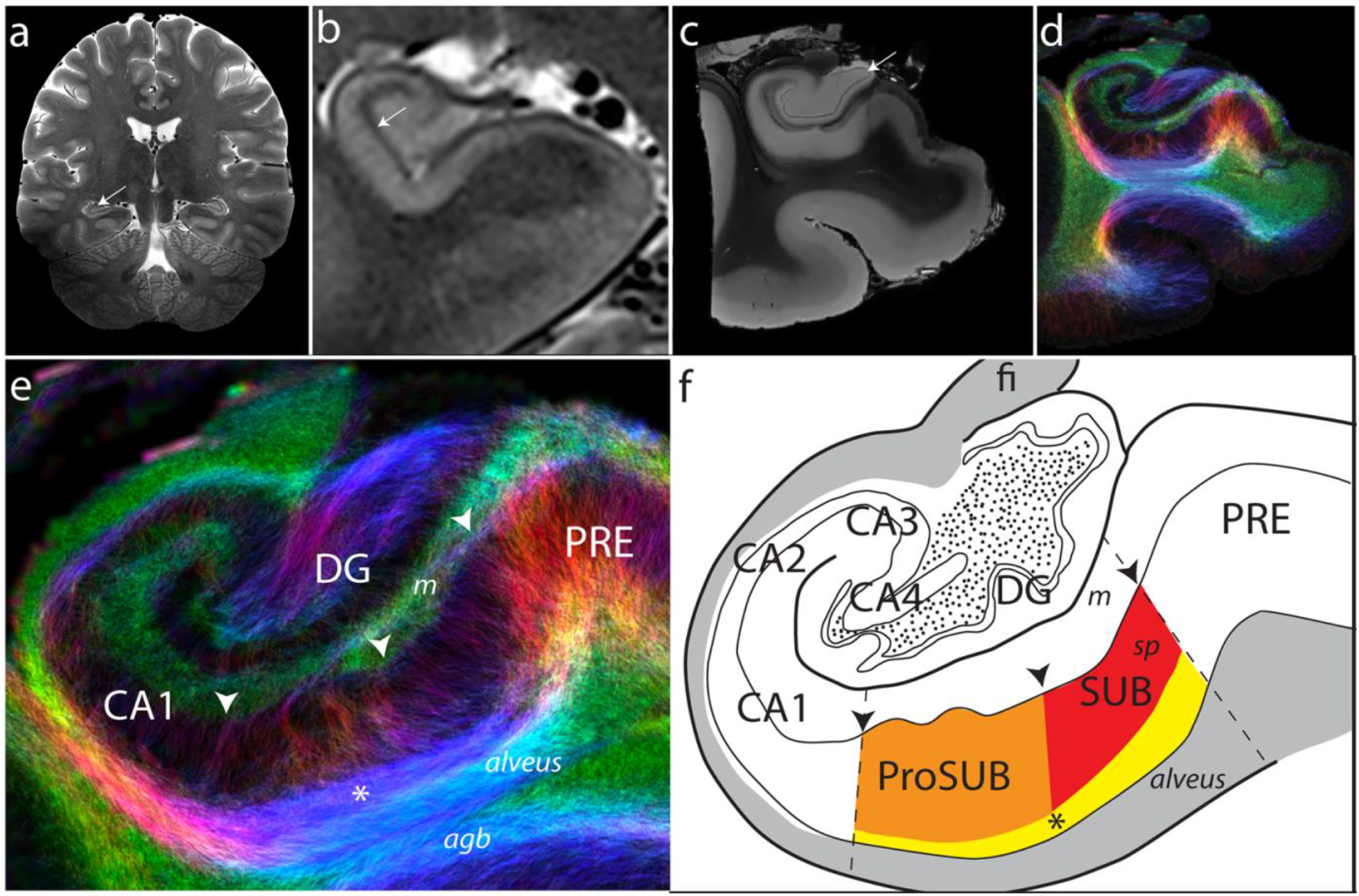
Tissue microstructure imaging resolution compared to gene expression segmentation. **a** 2D turbo spin echo T2w whole brain 7T *in vivo* MRI image acquired at 200 micrometer in-plane resolution with 2mm thickness (arrow points to hippocampus). **b** Zoomed in 7T MRI image of the hippocampus at a different rostrocaudal level showing the major hippocampal strata. The strata radiatum and stratum lacunosum moleculare in hippocampus proper as well as the SUB molecular layer appear as a dark band (arrow) that can help distinguish major hippocampal subregions. **c** *Ex vivo* T1w 16.4T MRI image ((50 μm)^3^ resolution) of a dissected post-mortem tissue sample from the temporal lobe. At this resolution, additional anatomical features are apparent including the granule cell layer of the dentate gyrus (arrow). **d** Fiber track density of the postmortem tissue sample obtained from high resolution angular diffusion imaging at 16.4T (150 μm 3D isotropic resolution). **e** Zoomed in view of the fiber track density in the hippocampus of the post-mortem sample with comparison to the gene expression-based atlas segmentation (**f**). Arrowheads in **e** and **f** mark the corresponding gene expression domain boundary positions. The ProSUB is distinguished from the adjacent SUB by the presence of several thicker dorsoventrally-oriented fiber bundles (orange/red boundary in atlas). In addition, a mediolateral fiber track dorsal to the alveus (asterisk) corresponds to the deep layer of cells located in both the ProSUB and SUB (yellow atlas area). Abbreviations: agranular bundle (agb), dentate gyrus (DG), fimbria (fi), molecular layer of the subiculum (m), presubiculum (PRE), prosubiculum (ProSUB), stratum pyramidale of the subiculum (sp), subiculum (SUB).

### Comparative analysis of the laminar organization in mouse and human SUB

In general, our observations of human SUB and ProSUB gene expression patterns are consistent with our previous analyses of gene expression patterns in the mouse SUB and we have maintained this color scheme in the human atlas drawings ^6^ (**Fig. 7**). The mouse HGEA defines 5 SUB subregions based on the representation and distribution of four distinct gene expression pyramidal sublayers. First, a dorsal and ventral part of dorsal subiculum (SUBdd and SUBdv) consists of a pyramidal layer with two gene expression sublayers (sublayers 1 and 4). The SUBdd most closely represents the area many studies refer to as the distal SUB, whereas ProSUB (sublayers 3 and 4) may closely correspond to proximal SUB in other studies. In contrast, the ventral subiculum (SUBv) has a trilaminar pyramidal layer (layers 2, 3, and 4). The thickness of these layers changes near the ventral tip of the subiculum (SUBvv) distinguishing this region from the SUBv. A major challenge for comparison analysis of mouse and human hippocampal gene expression data is the difference in orientation of the hippocampus between species. Because of the rotation of the longitudinal hippocampal axis across evolution, coronal mouse hippocampal sections are not in the same sectioning plane relative to the longitudinal axis as coronal human sections. Coronal hippocampus sections in humans would most directly relate to horizontal sections in the mouse and vice versa. However, several key aspects of SUB organization can be understood even comparing tissue that is sectioned at different angles relative to the longitudinal axis.

**Figure. 7.**
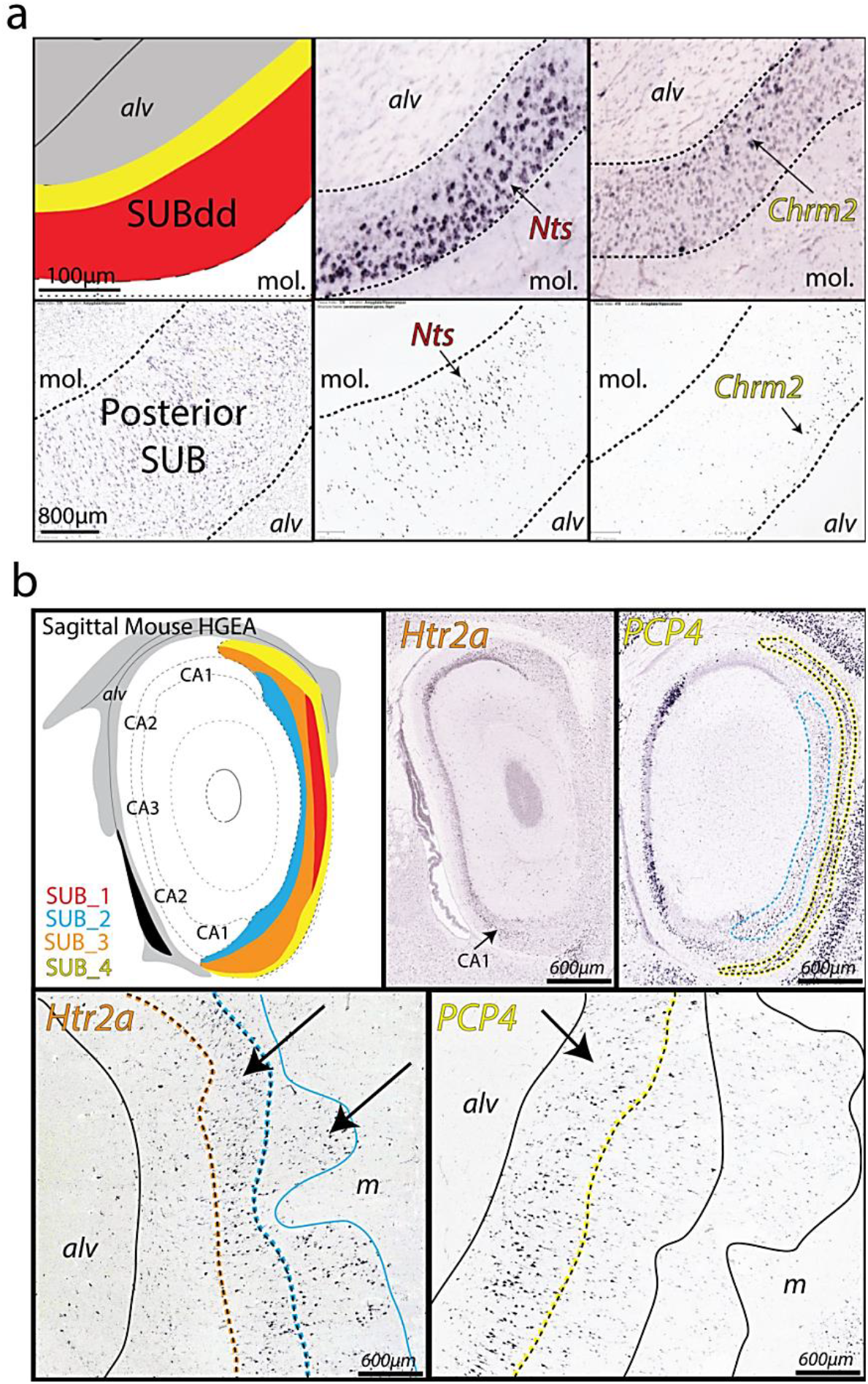
Homologous SUB laminar organization with similarities and differences in gene expression. **a** Comparison of *Nts* and *Chrm2* expression in the mouse dorsal subiculum (specifically SUBdd in the HGEA nomenclature, top row) and human posterior SUB (bottom row). Despite the difference in cell packing density, the laminar organization of the SUB pyramidal cells appears conserved across species. In the mouse HGEA, the *Nts*-expressing cells are located in SUB layer 1 (SUB_1), whereas *Chrm2* expression is located in SUB layer 4 (SUB_4). In both mouse and human, the *Nts*-expressing cells are located superficially near the molecular layer (m) whereas the *Chrm2*-expressing cells are located deep near the alveus (alv). The position of the alveus and molecular layer (as well as the laminar organization of the pyramidal layer) is dorsoventrally inverted due to the rotation of the hippocampal axis between the two species. **b** In contrast to the highly similar *Nts* and *Chrm2* expression patterns, *Htr2a* and *PCP4* are differentially expressed between the mouse (top, sagittal-cut images, HGEA atlas to left)) and human SUB (bottom, coronal sections). *Htr2a* expression is not strongly expressed in the mouse SUB (some minor expression in ventral CA1), but strongly expressed in human CA1 and SUB (bottom left). *PCP4’s* combinatorial expression pattern in SUB is bilaminar in the mouse (SUB_2 and SUB_4, top right) but only expressed in the deep layer of human SUB cells (corresponding to SUB_4 only, bottom right).

In both mouse dorsal SUB and human posterior SUB, *Nts* is strongly expressed in a superficial layer of SUB pyramidal neurons closest to the molecular layer, whereas *Chrm2*-expressing neurons are primarily located in the deepest part of the SUB, forming a continuous layer adjacent to the alveus white matter tract (**Fig. 7a**). Note that the position of the alveus and molecular layer are switched dorsoventrally in mouse vs. human tissue. This data suggests that the superficial SUB layer in humans (containing *Nts-* and *TH-*expressing neurons) is directly homologous to SUB sublayer 1 in the mouse HGEA, whereas the Chrm2-expressing deep layer in humans is homologous to mouse SUB sublayer 4. Therefore, *Htr2a*-expressing human SUB neurons likely correspond to the other two mouse SUB layers 2 and 3. However, *Htr2a* expression appears almost entirely absent in the mouse CA1 and SUB and other genes that are present in both mouse and human SUB (ex. *PCP4*) are expressed in different combinatorial patterns (**Fig. 7b**). Together, this data suggests that while the anatomical laminar organization is evolutionarily conserved between mice and humans, their exist both similarities and differences in the gene expression profile of the SUB layers and their unique cell types.

Although visualizing overall structure from thin-cut tissue sections can be difficult, the relationship of mouse and human SUB is easier to visualize in 3D and we have generated a model of the human SUB based on our understanding of the changes to the mouse SUB (**Fig. 8a**). In addition to the rotation of the longitudinal axis, one end of the hippocampus structure (the anterior pole in human corresponding to ventral pole in mouse) has folded back against the longitudinal axis. In comparison, the mouse and human SUB laminar organization demonstrate a similar subregional organization across the longitudinal axis despite its rotation (**Fig. 8b**). The area of the posterior human SUB containing gene expression layers 1, 3, and 4 correspond remarkably well to the SUBdd and ProSUB subregions in the dorsal part of the mouse SUB. In the anterior human SUB, the ventral area of SUB containing layers 1 and 4 is homologous to the mouse SUBdv whereas the dorsal part appears homologous to the trilaminar mouse SUBv/SUBvv. Additionally, the data from the human anterior pole is similar to the organization in the caudal parts of the mouse SUB where layers 1 and 4 partly curl around the ventral tip of the dentate gyrus. Overall, the data suggests the SUB has a conserved mammalian architecture that in humans has undergone expansion and structural folding.

**Figure. 8.**
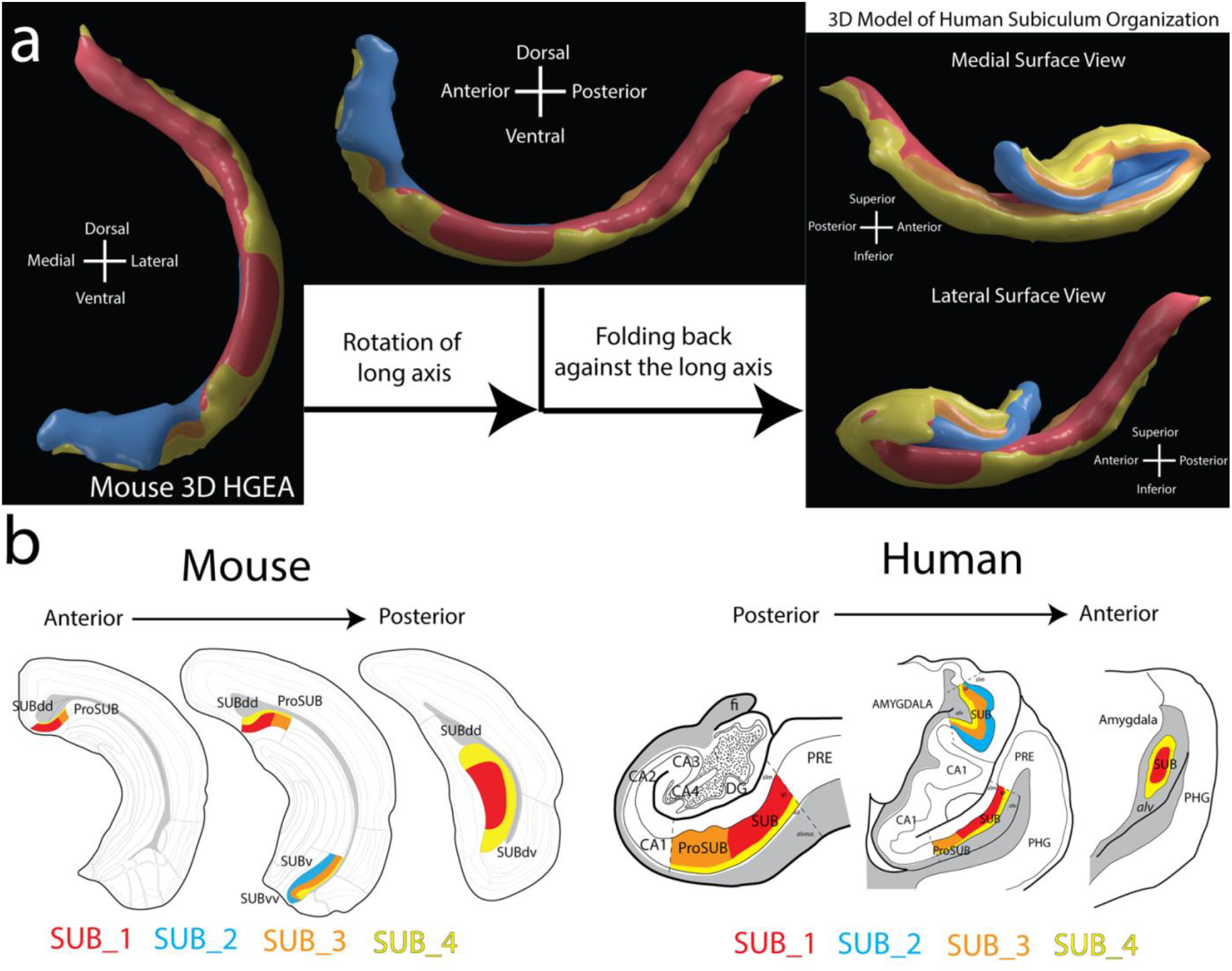
A 3D translational model of mouse and human SUB laminar organization. **a** Using the 3D HGEA model of the mouse SUB (left), we developed a 3D model of the human SUB based on our observations of the gene expression-based SUB lamina and the relative position of corresponding SUB subregions. Our model suggests two changes to the SUB have occurred across evolution between the mouse and human: 1) rotation of the longitudinal axis (middle), and 2) the folding back of the anterior SUB against the long axis (right). Images on the right show the view of the human SUB model from the medial (top) or lateral perspective (bottom; see also Supplementary Movie 2 or use Schol-AR app). **b** Coronal atlas section series from the mouse HGEA with the 4 colored gene expression layers and subregions (left) with similarly corresponding coronal human atlas drawings with similarly colored gene expression layers and subregions (right). Data viewable with Schol-AR augmented reality app, for details visit https://www.ini.usc.edu/scholar/download.html.

## Discussion

Overall, this study provides many novel insights into the organization of the human SUB and its evolutionary relationship to the mouse. Previous studies have analyzed gene expression patterns to map anatomical boundaries between the human SUB and ProSUB, however these studies did not report expression patterns of *Chrm2* and other genes which are present within a subset of neurons in both regions and demonstrate their laminar organization ^7–9^. Based on the current data, we believe that the structure of mouse and human SUB are highly conserved with respect to two major changes: 1) the rotation of the longitudinal axis and 2) the folding of the CA1/SUB at the anterior pole.

The mouse and human *in situ* hybridization data for *Nts*, *Chrm2*, and *Htr2a* suggest that the anatomical laminar organization of SUB pyramidal neuron cell types is conserved between mouse and human although many similarities and differences in their gene expression profiles can be observed. Generally, we find that the posterior SUB region in humans contains two distinct complementary gene expression layers (characterized by *Nts* and *Chrm2* expression) that suggests a homology to HGEA sublayers 1 and 4 found previously in the mouse SUBdd region. This gene expression data supports previous evidence for structural and functional homology between the posterior human SUB and dorsal mouse SUB and extends this relationship to the cellular level. Additionally, human sublayers 1 and 4 run continuously across the longitudinal axis as previously observed in mouse, suggesting that the anterior ventral part of the human SUB is homologous to the recently identified SUBdv mouse region ^6^. While sublayer 4 is present in the deepest part of the SUB at all levels, gene expression demarcating sublayer 1 is absent from the ProSUB and the more posterior parts of the dorsal anterior SUB near the amygdala. Instead, *Htr2a* was strongly expressed in superficial ProSUB neurons, similar in anatomical position to mouse HGEA sublayer 3 (although *Htr2a* is not expressed in mouse sublayer 3). Despite this difference, the overall *Htr2a* expression pattern in humans suggests an anatomical correspondence to HGEA sublayers 2 and 3 in mouse. In the dorsal anterior SUB region, the Nissl and gene expression data suggest three layers of cells: a deep *Chrm2*-expressing layer, an intermediate *Htr2a*-expressing layer with distinctive Nissl cytoarchitecture, and a superficial Htr2a-expressing layer. The trilaminar organization of the anterior dorsal SUB region (and its adjacent location to the amygdala) suggests a homology to the mouse HGEA SUBv/SUBvv regions which are composed of sublayers 2, 3, and 4. Although additional data is needed, the evidence presented here establishes a foundation for a translational HGEA map for describing mammalian hippocampal architecture across species.

The observed gene expression patterns highlight an important distinction between the dorsal and ventral parts of the human anterior SUB. The ventral part of the anterior SUB appears to be homologous to the SUBdv in mice whereas the dorsal part of the anterior SUB is homologous to the SUBv/SUBvv in mice. Based on the connectivity data in the mouse HGEA, this difference between the dorsal and ventral parts of the anterior human SUB implies differences in connectivity and function. If homologous to the mouse SUBv/SUBvv, then the neurons in the dorsal part of the anterior SUB would send output to limbic brain regions such as the amygdala, hypothalamus and prefrontal cortex. In contrast, the ventral part of the anterior SUB would be connected with the retrosplenial cortex and anterior thalamus and more related to visuospatial navigation (similar to SUBdv). In the future, human imaging studies of the SUB should separately analyze the dorsal and ventral parts of the anterior SUB to dissociate their anatomical/functional relationships.

Gene expression pattern analysis provides a useful way to map the anatomical distribution of distinct classes of neuronal cell types. Human imaging segmentation of the hippocampus and SUB are limited by the relatively low resolution of MRI contrast and the identification and position of hippocampal boundaries is highly inconsistent across studies ^22,23^. Due to the difficulty of drawing hippocampal boundaries on relatively homogenous MRI contrast, the current consensus among the community is to follow geometrical selection guidelines ^24–29^. However, a gene-expression based atlas could provide insight as to mechanistic and cell type-specific architecture of the hippocampus. Gene expression mapping provides a cellular resolution approach that could define ‘ground truth’ delineations and then be applied to MRI datasets. In support of this approach, we have found evidence that the gene expression boundaries reflect anatomical differences in the microstructure of human tissue (**Fig. 6**). Although it is currently unclear if these differences in fiber orientation reflect differences in long range connectivity or cell type-specific morphologic differences, our previous study found a strong relationship between anatomical connectivity and gene expression within the mouse hippocampus ^6^. Notably, our previous study found that mouse sublayer 4 neurons had a high degree of interconnectivity along the transverse hippocampal axis and the fiber tract identified in the TDI fiber analysis suggests a similar connectivity pathway exists in humans. As the resolution of MRI imaging technology continues to improve and microstructural imaging becomes clinically feasible and quantitatively-specific ^30–32^, we believe the data here provides an important foundation for an integrative approach to bridge the gap between MRI imaging and human tissue histology.

### Considerations for investigations of translational mammalian neuronal cell types

Recently, a translational gene expression study comparing mouse and human suggested a conservation of cortical cell types with divergent features across mammals ^33^. In the hippocampus, *CALB1* has previously been shown to be selectively expressed in the human dentate gyrus (DG), whereas CALB1 in the mouse is expressed within the DG, CA1, and CA2 ^34^. These variations raise important considerations as to how to classify cell types across species: Do the CALB1-positive CA1 and CA2 neurons represent a distinctly murine neuronal cell type not found in humans or are the homologous cell types still present in the human, but simply no longer express CALB1? Although the current study focused on the SUB, we also observed several instances where the combinatorial expression of specific genes is different between mouse and human across a variety of hippocampal regions. Cross-species differences at the single-cell transcriptome level may underlie the failure to successfully translate mouse research to the clinic. Drugs that are effective in mice because they bind a receptor in specific target neurons may not work in humans because 1) either the homologous human neuron doesn’t express that receptor or 2) the receptor is expressed in additional neurons causing unexpected side effects that were not observed in the mouse experiments. Because of these potential pitfalls in cross species drug development, it is critical that we understand the homologous single-cell gene expression profiles between mouse and human neuronal cell types. Ultimately, a convergence of data modalities describing anatomical, molecular, and physiological characteristics is imperative to accurately defining cell types across different mammalian nervous systems. The evidence presented here provides a foundation for understanding the organization of mouse and human subiculum, but further characterization is necessary for a comprehensive map of spatial gene expression patterns similar to the mouse HGEA ^6^.

### Conclusions

This study establishes a new understanding of the organization of the human SUB and provides a translational perspective that corroborates and extends the anatomical and functional homology across mouse and human SUB. We find that both mouse and human SUB pyramidal neurons contain a hidden laminar organization of gene expression across the longitudinal axis. These findings establish a foundation for future investigations of the organization of the human hippocampus using gene expression and establishing an integrative approach with *in vivo* and *ex vivo* MRI imaging.

## Methods

### Allen Brain Atlas in situ hybridization data

All human and mouse *in situ* hybridization images used in the analysis of this study were downloaded from the Allen Brain Atlas website (www.brain-map.org). Complete detailed information about histological processing and hybridization can be found in the Allen Institute white paper (Mouse: “*In situ* hybridization data production”, Nov. 2011; http://help.brain-map.org/display/mousebrain/Documentation; Human: “*In Situ* Hybridization in the Human Brian Atlas”, Oct. 2013 v.7; http://help.brain-map.org/display/humanbrain/Documentation).

Briefly, human *in situ* hybridization image datasets from a variety of brain regions are categorized according to five specific projects (Cortex Study, Neurotransmitter Study, Subcortex Study, Schizophrenia Study, and Autism Study). Only the Neurotransmitter Study currently contains hippocampus/amygdala tissue samples. In the Neurotransmitter Study, the right hemisphere of frozen brains from four post-mortem subjects (H0351.1009, a 57 year old Caucasian male with hypertension; H0351.1010, 28 year old Hispanic male control; H0351.1012, 31 year Caucasian male control; H0351.1016, 55 year old Caucasian male control) were sectioned into thick 1-1.5cm coronal slabs (‘control’ tissue from patients with no known neuropsychiatric disease, see www.brain-map.org for additional subject information). Coronal slabs were frozen in a bath of dry ice and isopentane and stored at −80°C until sectioning.

Prior to sectioning, coronal slabs were blocked into samples measuring approximately 4.2cm × 3.7cm to capture the primary brain structures of interest. Frozen tissue samples were sectioned coronally, from anterior to posterior, at 20μm thickness using a Leica CM3050 S cryostat. For the Neurotransmitter Study, a set of *in situ* hybridization digoxigenin-labeled probes for 88 genes were designed to assay genes regulating a variety of neurotransmitter systems across the brain so not all genes are expressed within hippocampus (Gene list: *Abat, Ache, Adcyap1, adora2a, Aldh2, bdnf, calb1, calb2, cartpt, cbln2, chat, chrm2, chrm3, chrna2, chrna3, chrna7, chrnb2, cnr1, cnrip1, crym, dld, drd2, gabra1, gabra2, gabra3, gabra4, gabra5, gabrb1, gabrb2, gabrb3, gabrd, gabre, gabrg1, gabrg2, gabrq, gad1, gad2, gfap, glra1, glra3, glrb, gls, got1, got2, gria1, gria2, gria4, grik1, grik2, grin1, grin2a, grin2b, grin3a, grm1, htr1a, htr2a, htr2c, maob, mfge8*, mbp, nefh, ngb, nnat, npy1r, nts*, nxph1, oxtr, pcp4, pdyn, penk, pnoc, pvalb, scg2, slc17a6, slc17a8, slc1a1, slc1a2, slc1a3, scl1a4, slc32a1, slc6a1, syt2, tac1, tac3, th, trh, trhr*,* asterisked gene data not available from all subjects). Horseradish peroxidase conjugated antibody based colorimetric reaction of the digoxigenin-probes produces a blue/purple precipitate within neuronal cell bodies containing targeted mRNA transcripts (see Allen Institute white paper for complete details; http://help.brain-map.org/display/humanbrain/Documentation). All 88 probes and additional histological stains are performed in sequential tissue sections in order from anterior to posterior (Nissl stained sections occur every 25 sections). Each tissue section is hybridized to one unique probe and the procedure is repeated every 100 sections such that the rostrocaudal sequence of each gene expression tissue series is spaced 2mm apart (100 sections × 20μm section thickness). All image data used for analysis was openly published online and downloaded from www.brain-map.org.

### Post-mortem tissue and ex vivo imaging parameters

The hippocampal specimens used for *ex vivo* imaging was obtained post-mortem from a 55-year-old male with normal cognitive function at the Keck Medical Center of USC (Los Angeles, CA). The specimens were immediately fixed in 10% phosphate-buffered formalin, and later dissected at the level of the lateral geniculate nucleus, including the hippocampal and parahippocampal gyri. A 1cm extent of the hippocampal tissue was submitted for *ex vivo* MRI analysis.

MRI data was acquired on a 16.4T vertical wide-bore microimaging system, running Paravision 6.0.1 (Bruker Biospin, Karlsruhe, Germany), using a micro 2.5 gradient coil (max strength 1.5T/m) and 28 mm birdcage volume coil (M2M Imaging, Brisbane, Australia). The sample was incubated in 0.2% gadolinium diethylenetriaminepentaacetic acid (Gd-DTPA, Magnevist, Bayer) for 4 days, and fixed onto a plastic holder with a small amount of cyanoacrylic glue. To reduce geometrical distortion and preserve the sample during MRI, it was immersed inside a polyperfluoroether medium (Fomblin Y06/06, Solvay Solexis, Italy) ^32,35,36^.

DW-MRI was acquired using 3D Stejskal-Tanner ^37^ spin-echo sequence to achieve high signal-to-noise ratio (SNR) at (150 μm)^3^ isotropic resolution. Data was acquired with three diffusion weightings: b-values of 1000, 3000 and 5000 s/mm^2^, with 20, 30 and 45 diffusion encoding gradient directions with distinct optimized spherically even distribution ^38^. In addition, a total of 6 unweighted images (B0) were acquired. 3D DW-MRI spin-echo was acquired using FOV = 25.5 × 20 × 23.4 cm and matrix size = 170 × 130 × 156. TE, pulse duration and separation times were fixed across shells to avoid time-dependent effect on diffusion signal. Total DW-MRI scan time was 48 hours and 22 minutes (also at 22°C).

### *Ex vivo* MRI data analysis

Image volumes were corrected for Gibbs ringing (using Trapezoid windowing) and N4 field bias (using ANTs software: http://stnava.github.io/ANTs/). DW-MRI data quality and signal values were visually inspected for quality control. Track-density imaging (TDI) ^39^ was performed using single shell DW-MRI on the shell with b-value of 5000 s/mm^2^, using MRtrix software (version 0.2.12; http://jdtournier.github.io/mrtrix-0.2/index.html). Voxels with FA>0.7 were segmented and the spherical harmonic decompositions of all the resulting profiles were then averaged to estimate the response function. We then applied constrained spherical deconvolution ^40^ to estimate the fiber orientation distribution in each voxel using a maximum spherical harmonic of order 6. Then, 500,000 streamlines were generated using probabilistic tractography tool ^41^ with the following parameters: curvature=0.075, cutoff=0.1, minlength=1, length=15, step=0.015. TDI with voxel size of (100 mm)^3^ was then derived from generated streamlines.

### *In vivo* high resolution hippocampal imaging

A 32-year-old female was scanned using T2-weighted turbo spin echo sequences with 2mm slice thicknesses and 340 µm (interpolated to 170 µm) in-plane resolution, 4 averages, resulting to total scan time of 13 minutes ^42,43^. We used a single-channel quadrature transmit radiofrequency (RF) coil and a 32-channel receive array coil (Nova Medical Inc., MA). The institutional review board of the University of Southern California approved the study. Informed consent was obtained from the volunteer, and the image was anonymized.

## Acknowledgments

This work was supported by the National Institute of Mental Health grants R01 MH094360-01A1 (H.-W.D.), U01 MH114829-01 (H.-W.D.), and RF1 MH114112-01 (H.-W.D) and National Institute of Neurological Disorders and Stroke grant R21NS091586-02 (C.A.M.). We acknowledge the supports from the Queensland NMR Network and the National Imaging Facility (a National Collaborative Research Infrastructure Strategy capability) for the operation for 16.4T MRI at the Centre for Advanced Imaging, the University of Queensland.

